# A global indicator of the capacity of terrestrial ecosystems to retain biological diversity under climate change: the Bioclimatic Ecosystem Resilience Index

**DOI:** 10.1101/795377

**Authors:** Simon Ferrier, Thomas D Harwood, Chris Ware, Andrew J Hoskins

**Affiliations:** CSIRO Land and Water, GPO Box 1700, Canberra ACT 2601, Australia; CSIRO Health and Biosecurity, James Cook Drive, Townsville 4811, Australia

**Keywords:** biodiversity, indicator, ecosystem resilience, climate change, terrestrial, global

## Abstract

An important element of the Convention on Biological Diversity’s Aichi Target 15 – i.e. to enhance “ecosystem resilience … through conservation and restoration” – remains largely unaddressed by existing indicators. We here develop an indicator addressing just one of many possible dimensions of ecosystem resilience, by focusing on the capacity of ecosystems to retain biological diversity in the face of ongoing, and uncertain, climate change. The Bioclimatic Ecosystem Resilience Index (BERI) assesses the extent to which a given spatial configuration of natural habitat will promote or hinder climate-induced shifts in biological distributions. The approach uses existing global modelling of spatial turnover in species composition within three broad biological groups (plants, invertebrates and vertebrates) to scale projected changes in composition under a plausible range of climate scenarios. These projections serve as filters through which to analyse the configuration of habitat observed at a given point in time (e.g. for a particular year) – represented as a grid in which cells are scored in terms of habitat condition. BERI is then calculated, for each cell in this grid, as a function of the connectedness of that cell to areas of natural habitat in the surrounding landscape which are projected to support a similar composition of species under climate change to that currently associated with the focal cell. All analyses are performed at 30-arcsecond grid resolution (approximately 1km cells at the equator). Results can then be aggregated to report on status and trends for any desired set of reporting units – e.g. ecoregions, countries, or ecosystem types. We present example outputs for the Moist Tropical Forest Biome, based on a habitat-condition time series derived from the Global Forest Change dataset. We also describe how BERI is now being extended to cover all biomes (forest and non-forest) across the entire terrestrial surface of the planet.

## 1. Introduction

In 2010 the Conference of the Parties of the Convention on Biological Diversity (CBD) adopted a Strategic Plan for Biodiversity for the period 2011 to 2020 (SCBD 2010). This plan includes 20 ambitious targets – collectively known as the Aichi Targets – nested under five Strategic Goals addressing underlying causes of biodiversity loss, reducing direct pressures, improving the status of biodiversity, enhancing benefits from biodiversity and ecosystem services, and enhancing implementation through appropriate planning, management and capacity building. A wide range of indicators have been developed, and are already being used, to assess progress towards achieving the Aichi Targets (Tittensor et al. 2014). However, not all of the targets are well supported by effective and readily available indicators. A recent review of gaps in the coverage of indicators by the Biodiversity Indicators Partnership (a global initiative coordinating the development and delivery of many of the indicators used by the CBD) has identified three Aichi Targets which are particularly poorly served by their existing indicator suite (McOwen et al. 2016). One of these is Target 15: *“By 2020, ecosystem resilience and the contribution of biodiversity to carbon stocks have been enhanced, through conservation and restoration, including restoration of at least 15 per cent of degraded ecosystems, thereby contributing to climate change mitigation and adaptation and to combating desertification.”*

As noted by McOwen et al (2016), any attempt to assess progress against Target 15 is challenged by the broad and multi-faceted coverage of this target, along with the vagueness with which different elements of the target are defined, leaving these elements open to diverse interpretations. Some elements of Target 15 are potentially amenable to assessment using indicators already developed, or being developed, in other policy domains. For example, global indicators of carbon stock changes are well-established under the auspices of the United Nations Framework Convention on Climate Change (e.g. Petrescu et al. 2012), and a new composite indicator of land degradation is currently being developed under the auspices of the United Nations Convention to Combat Desertification (Sims et al. 2017).

One element of Target 15 which remains largely unaddressed by existing indicators is that relating to the enhancement of “ecosystem resilience”. This element, perhaps more than any other included in the target’s formulation, is left open to a wide range of possible interpretations. We assume here that the broad definition of resilience of most relevance in this particular context (amongst alternatives such as engineering resilience and social-ecological resilience) is that of ecological resilience – i.e. *“the ability of a system to withstand shock and maintain critical relationships and functions”* (Quinlan et al. 2016). However, even accepting this definition, developing indicators to assess change in the resilience of whole ecosystems is challenged by the reality that, as observed by Walker and Salt (2012) *“resilience is not a single number or a result. It is an emergent property that applies in different ways and in the different domains that make up your system. It is contextual and it depends on which part of the system you are looking at and what questions you are asking”*. This means that no single indicator will be able to address all possible dimensions of ecosystem resilience (Hodgson et al. 2015). Different indicators will therefore be needed to address different answers to the question *“resilience of what to what?”* (Carpenter et al. 2001). In other words, it is vital that we clearly define what any indicator developed to assess change in ecosystem resilience is actually measuring – both in terms of the component or attribute of the ecosystem we are interested in maintaining (i.e. resilience “of what?”) and in terms of the disturbance or pressure we want the system to withstand in maintaining this component (i.e. resilience “to what?”).

The indicator we describe in this paper focuses on just one aspect of ecosystem resilience – the capacity of ecosystems to retain biological diversity in the face of ongoing, and uncertain, climate change. This particular focus aligns well Target 15’s framing of ecosystem resilience as “contributing to climate change mitigation and adaptation”. It also aligns strongly with rapid growth in awareness over recent years of the need to better consider potential impacts of climate change on biodiversity, in assessment and planning efforts around ecosystem conservation and restoration (Groves et al. 2012; Stein et al. 2013).

We here view the effects of ecosystem degradation, conservation and restoration on the capacity of ecosystems to retain biological diversity under climate change as playing out across two broadly different scales, corresponding with the concepts of “local system resilience” and “spatial resilience” (Cumming 2011). At the local scale, ecosystem degradation, conservation and restoration will affect the capacity of a given location to support species native to the ecosystem concerned, largely through effects on the overall condition of habitat for these species. Enhancing habitat condition at a location is also expected, in turn, to enhance capacity to accommodate potential change in the composition of supported species under climate change (Eigenbrod et al. 2015). If this were the only determinant of the capacity of ecosystems to retain biological diversity under climate change then an indicator of this capacity could be derived simply and directly from mapped changes in habitat condition or intactness across the planet (e.g. Newbold et al. 2016; Venter et al. 2016). However such an approach would not consider the potential need for species to shift their distributions in response to changing climatic conditions (Pecl et al. 2017). This requires consideration of an important manifestation of spatial resilience. The capacity of ecosystems to accommodate climate-induced shifts in species distributions will be a function not only of the mean condition of habitat across a landscape or region of interest, but also of the configuration and connectedness of that habitat in both geographical and environmental space (McGuire et al. 2016).

To integrate consideration of both local and spatial resilience in the indicator developed here we start by assuming availability of a spatial grid for the region of interest, in which the value for each grid-cell is a best estimate of the condition of habitat within that cell at a given point in time (e.g. for a particular year of observation). We also assume that the cumulative effects of ecosystem degradation, or conversely of ecosystem conservation and restoration, up to this point in time will be manifested in this grid. Likewise, if a time series of such grids is available (e.g. for different years of observation), then this is assumed to provide a reasonable estimate of changes in habitat condition over time, and to thereby serve as a reasonable foundation for assessing change, over this same time period, in the capacity of ecosystems to retain biological diversity in the face of climate change. The challenge we focus on in this paper is therefore developing an effective, yet practical, means of building consideration of spatial resilience on top of this foundation.

How can we assess the extent to which a given spatially-explicit configuration of habitat condition within a given region confers capacity to retain biological diversity by allowing species’ distributions to shift, if necessary, in response to climate change? Two broad analytical approaches have dominated efforts over the past 20 years to assess potential impacts of climate change on the spatial distribution of biodiversity. The first, and most widely applied, approach focuses on modelling shifts in the distribution of particular species as a spatially explicit function of projected change in climate (Elith and Leathwick 2009). The popularity of this approach has resulted in a proliferation of correlative and mechanistic modelling tools, and of techniques for addressing a growing number of relevant biological and ecological factors within these models – e.g. dispersal capacity, population dynamics, evolutionary adaption, lag effects, species interactions (Urban et al. 2016).

Concerns regarding sparseness and unevenness in the geographic and taxonomic coverage of data needed for such modelling, and the high level of uncertainty associated with many aspects of modelled responses to climate change, have at times motivated interest in an alternative analytical approach. This involves focusing purely on analysing spatiotemporal patterns in abiotic environmental variables alone, without any direct use of biological data or any explicit modelling of biological responses. The manifestation of this approach attracting most interest in recent years is the analysis of climate velocity – i.e. the speed at which a hypothetical species associated with a given geographic location would need to move to maintain its current climatic conditions under climate change (Brito-Morales et al. in press). Along similar lines, interest has been growing rapidly around adaptation strategies aiming to enhance the capacity of landscapes to retain biological diversity by “conserving nature’s stage” (Beier et al. 2015) – i.e. ensuring that conserved areas encompass high levels of diversity in, and therefore steep gradients of, abiotic attributes including climate (Heller et al. 2015), but also considering topography and soils (Schloss et al. 2011).

An arguable strength of this second approach – i.e. working with environmental data alone – is its utility for addressing regions and/or components of biodiversity where the data and understanding required to explicitly model biological responses are lacking. An important limitation, however, lies in the implicit assumption that a given magnitude of spatial and/or temporal change in a climate variable, or any other environmental variable for that matter, can serve as a reasonable proxy for the amount of biological change expected. Many decades of ecological and biogeographic research suggest cause for concern regarding this assumption. Even under present-day conditions, the level of turnover in biological composition observed between two locations exhibiting a given difference in a climatic attribute – e.g. a 2°C difference in mean annual temperature – can vary dramatically depending on the biological group of interest, the climatic position of the locations (e.g. at the low versus high end of a temperature gradient), and the biogeographic history of the region concerned (Buckley and Jetz 2008; Fitzpatrick et al. 2013; Konig et al. 2017).

To accommodate these sources of variation in developing the indicator described here, we adopt a third alternative for assessing the capacity of landscapes to retain biological diversity under climate change which effectively occupies the middle ground between explicit modelling of shifts in biological distributions, and analyses based on spatiotemporal patterns in climate alone. This approach uses best-available location records for large numbers of species, combined with statistical modelling of spatial turnover in species composition, to scale (transform) multidimensional environmental space, such that distances within this transformed space correlate as closely as possible with observed levels of biological turnover (Ferrier et al. 2007). Through space-for-time substitution, we use this scaling to assess the extent to which an observed configuration of habitat condition (e.g. for a given region in a given year) will promote or hinder the connectivity of shifting environments, under a plausible range of climate futures.

The general approach underpinning development of the indicator described here has been applied to a wide variety of assessments of biodiversity conservation status, climate-change vulnerability, and adaptation options, for the Australian continent over the past 10 years (e.g. Bryan et al. 2014a; Ferrier et al. 2012; Prober et al. 2012; Williams et al. 2015). Recent expansion of this analytical capability to cover the entire terrestrial surface of the planet (Hoskins et al. 2019) has already facilitated the establishment of three global indicators for assessing progress against Aichi Targets 5 and 11 (GEO BON 2015) – the Biodiversity Habitat Index (BHI; https://www.bipindicators.net/indicators/biodiversity-habitat-index) and the Protected Area Representativeness and Connectedness (PARC) Indices (https://www.bipindicators.net/indicators/protected-area-representativeness-index-parc-representativeness; https://www.bipindicators.net/indicators/protected-area-connectedness-index-parc-connectedness). We here add a new indicator to this suite – the Bioclimatic Ecosystem Resilience Index (BERI) – tailored specifically to assess change in the capacity of ecosystems to retain biological diversity under climate change, thereby addressing a key aspect of “ecosystem resilience” under Aichi Target 15.

## 2. Material and methods

### 2.1 General framework

The overall methodological framework for deriving the BERI indicator is depicted in Figure 1. This framework is underpinned by global models of spatial turnover in species composition previously established as part of CSIRO’s Biogeographic modelling Infrastructure for Large-scale Biodiversity Indicators (BILBI; Hoskins et al. 2019). These correlative models predict the compositional turnover (also known as “compositional dissimilarity” or “pairwise beta diversity”) expected between any two grid-cells on the planet as a function of fine-scaled spatial variation in climate, terrain and soils within major biomes and biogeographic realms. The models were fitted to best-available occurrence records for large numbers of species, and best-available environmental surfaces, using generalised dissimilarity modelling (GDM; Ferrier et al. 2007). Nonlinear functions generated by this model-fitting process describe the relative importance of different environmental gradients in driving spatial turnover in species composition, and how rates of turnover vary between different positions along each of these gradients.

**Figure 1:**
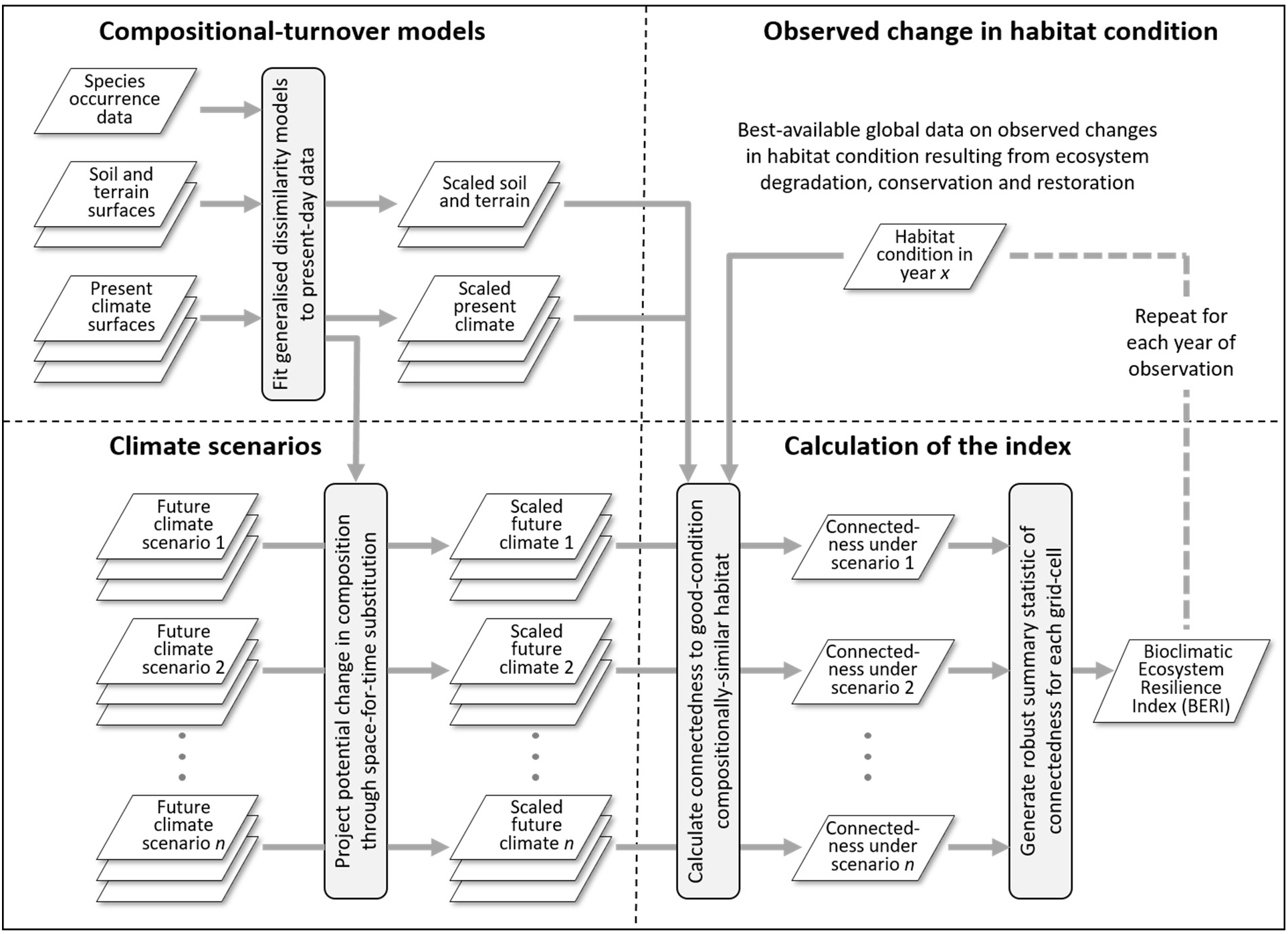
Overall methodological framework for deriving the BERI indicator

For the purposes of deriving the BERI indicator, these functions are used to project potential shifts in species composition over time, under a plausible range of climate scenarios. These projected shifts in species composition then serve as a set of filters through which to assess the capacity of an observed configuration of habitat condition to retain biological diversity under climate change. This capacity is assessed in relation to each individual grid-cell in turn (serving as a “focal cell”) by analysing the connectedness of this cell to areas of habitat in the surrounding landscape which are both: 1) in relatively good condition; and 2) projected to support an assemblage of species under plausible future climates which is similar in composition to that associated with the focal cell under present climatic conditions. The value of BERI assigned to each cell is then calculated as a robust summary statistic of resulting levels of connectedness for the evaluated climate scenarios. This is expressed as a ratio relative to the connectedness achievable if the cell were surrounded by a continuous expanse of pristine habitat, and were not subjected to any change in climate.

### 2.2 Compositional-turnover models

The compositional-turnover models we use here to derive the BERI indicator have been described in detail by Hoskins et al. (2019). These GDM models were fitted to available occurrence records for all terrestrial species within the following taxa: vascular plants, amphibians, reptiles, birds, mammals, ants, bees, beetles, bugs, butterflies, centipedes, dragonflies, flies, grasshoppers, millipedes, snails, moths, spiders, termites, and wasps. The records for amphibians, birds and mammals were extracted from data accessible through the Map of Life (https://mol.org/) while records for all other taxa were extracted from data accessible through the Global Biodiversity Information Facility (GBIF; http://www.gbif.org/).

A total of 183 separate GDM models were fitted, one for each possible combination of three broad biological groups – vascular plants, vertebrates and invertebrates – and 61 bio-realms (unique combinations of biomes and biogeographic realms, as per the World Wildlife Fund (WWF) global classification of terrestrial ecoregions (Olson et al. 2001). The total global datasets used to fit these models consisted of: 52,489,096 records of 254,145 species of vascular plants; 33,549,534 records of 24,442 species of vertebrates; and 13,244,784 records of 132,761 species of invertebrates. To accommodate the presence-only nature of much of the assembled biological data, GDM models were fitted to observed matches and mismatches in species identity between pairs of individual occurrence records (see Hoskins et al. 2018 for details).

The environmental surfaces used to fit the GDM models were at 30-arcsecond grid resolution (a cell size of approximately 1km x 1km at the equator), and therefore all predictions of compositional turnover from the models are at this same resolution. Climatic predictors of compositional turnover were derived from long-term average climate surfaces accessible through WorldClim (http://www.worldclim.org/)(Hijmans et al. 2005) and included: minimum temperature of the coolest month; maximum temperature of the warmest month; maximum monthly diurnal temperature range; annual precipitation; annual actual evaporation; potential evaporation of driest month; maximum monthly water deficit; and minimum monthly water deficit. The temperature, evaporation and water-deficit variables were also adjusted for the radiative-shading effects of topography. For details of this adjustment, and for descriptions of soil and terrain variables included alongside climate in the modelling, see Hoskins et al. (2019).

### 2.3 Climate scenarios

We use the fitted GDM functions, describing rates of turnover in present-day species composition along each of the environmental gradients, to project potential levels of compositional turnover expected under plausible future scenarios of climate change. This is undertaken using standard space-for-time substitution – i.e. the same fitted model used to predict compositional turnover between two grid-cells under present-day environmental conditions is here used to predict turnover between a cell under present conditions and either itself, or another cell, under the future conditions expected for a given climate scenario (for further explanation of this approach see Blois et al. 2013; Ferrier et al. 2012; Fitzpatrick et al. 2011).

We first generated projected grids for the same eight climatic predictors, used to fit the present-day GDMs, from downscaled CMIP5 climate projections accessible through WorldClim (http://www.worldclim.org/CMIP5v1), again adjusting these for the radiative-shading effects of topography. Each of these grids was then scaled (transformed) using the relevant GDM function describing how rates of turnover in present-day species composition vary along that particular gradient, for a given biological group within a given bio-realm. While computationally demanding, this scaling of future climate grids needed to be performed only once. The resulting GDM-scaled climate grids allow very efficient prediction of compositional-turnover values across time and space when calculating the BERI indicator repeatedly for different configurations of habitat condition (see next subsection).

We have generated these scaled grids for six climate scenarios, all projected to 2050. Each of these scenarios combines a particular climate model (general circulation model; GCM) with a particular level of greenhouse gas concentration (representative concentration pathway; RCP). Four of the scenarios employ the IPSL-CM5A-LR GCM, combined with RCP 2.6, RCP 4.5, RCP 6.0 and RCP 8.5 respectively. These scenarios align directly with those employed in a major inter-comparison of biodiversity and ecosystem-service models recently undertaken by the Intergovernmental Platform for Biodiversity and Ecosystem Services (IPBES) Expert Group on Scenarios and Models (Kim et al. 2018). We adopt them here to account for uncertainty in future climate trajectories associated with potential variation in greenhouse gas emissions. To account for potential differences between climate models we have further combined RCP 8.5 (the highest greenhouse-gas concentration) with two alternative GCMs: ACCESS1-0 and GFDL-CM3.

### 2.4 Calculation of the index

The BERI index can be calculated for any 30-arcsecond grid of habitat condition observed at a given point in time (e.g. for a particular year). Each cell in this grid is assumed to contain a value between 0, equating to complete local loss of native species, and 1, equating to complete local retention of native species (potential sources of such data are considered in the following sub-section). The index is initially calculated in relation to each individual cell – i.e. every cell serves, in turn, as the focal cell for assessing connectedness to areas of good-condition habitat in the surrounding landscape which are projected to support a similar composition of species under climate change to that currently associated with this cell. Once BERI values have been generated at the grid-cell level, summary statistics for larger reporting units (e.g. countries, ecoregions) can be derived simply by averaging the values of all cells encompassed by a given unit.

Connectedness is assessed using an extension of the cost-benefit approach (CBA) developed by Drielsma et al (2007) which is founded on well-established principles of meta-population ecology. In the interests of computational efficiency, we adapt Drielsma et al’s use of irregular “petals” to instead divide the neighbourhood around each focal cell into 80 segments (Figure 2), by intersecting eight radial sectors (N, NE, E, SE, S, SW, W, NW) with 10 radial rings (with outer edges at 2, 5, 10, 25, 50, 100, 200, 300, 400, and 500km). The effective distance *d*_*ij*_ between focal cell *i* and segment *j* is estimated by finding the least-cost path connecting these two elements, traversing any set of segments falling within the radial sector containing segment *j* or the sectors immediately adjacent to this. The contribution that any intervening segment makes to *d*_*ij*_ is calculated as the map-distance traversed across that segment divided by the mean habitat-condition of all 30-arcsecond cells contained within the segment.

**Figure 2:**
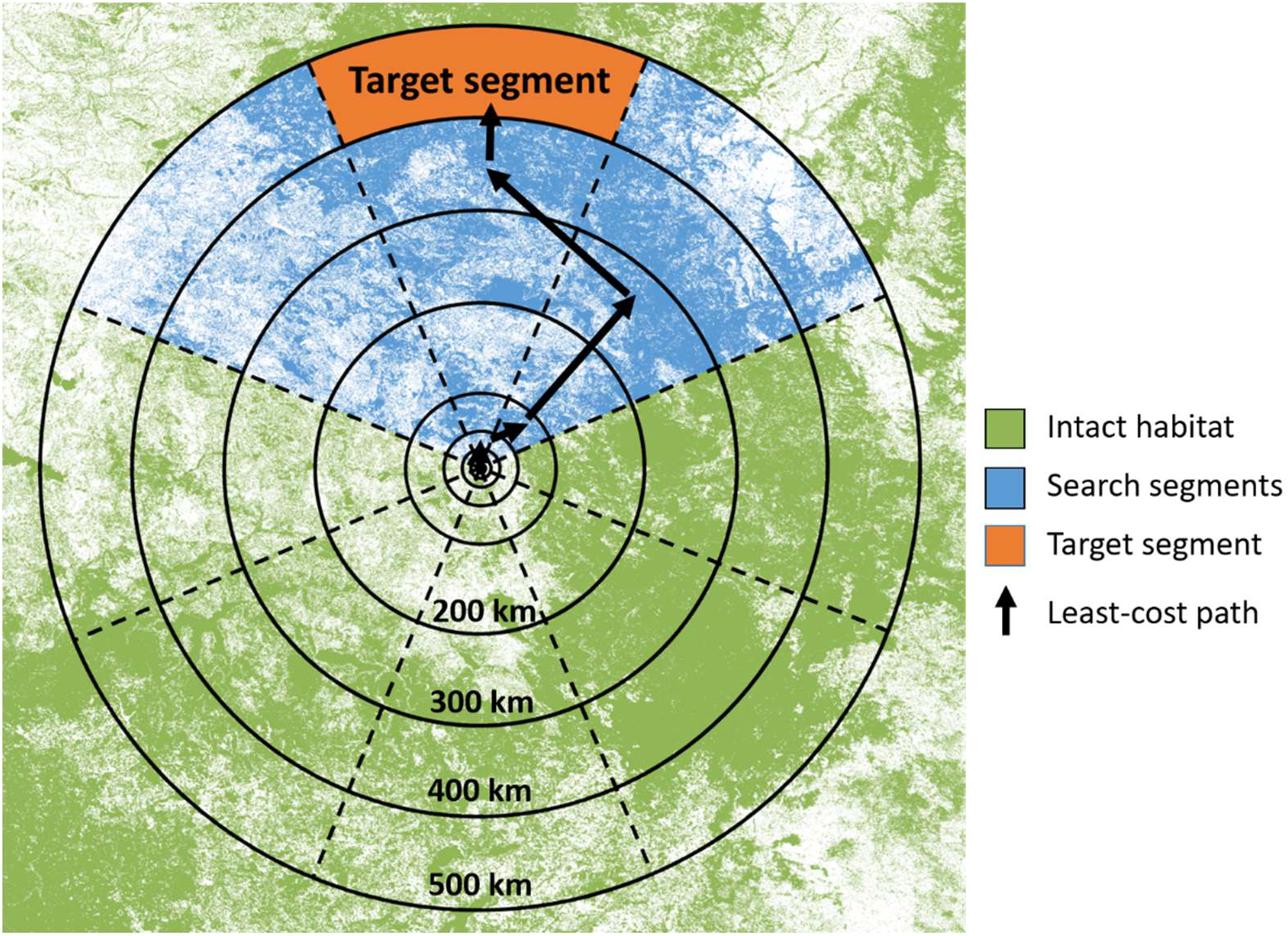
Radial sectors and rings dividing the neighbourhood around each focal cell into the 80 segments used in estimating connectedness.

The connectedness of focal cell *i* under climate scenario *k* (see Figure 3, panel d) to all segments is then calculated as:

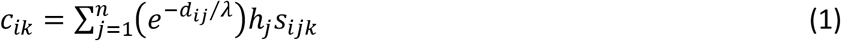

where *n* is the number of segments in the neighbourhood around the focal cell (80 in this case); *λ* is the median distance that species in the biological group of interest are expected to be able to disperse through intact habitat within the time period of the scenario, used to scale probability of dispersal as a negative exponential function of effective distance; *h*_*j*_ is the mean habitat-condition of all 30-arcsecond cells contained within segment *j*; and *S*_*ijk*_ is the mean compositional similarity predicted between the focal cell under present-day climate, and each of the cells in segment *j* under climate scenario *k*, assuming hypothetically that all these cells are in perfect habitat condition (see Figure 3, panel b).

**Figure 3:**
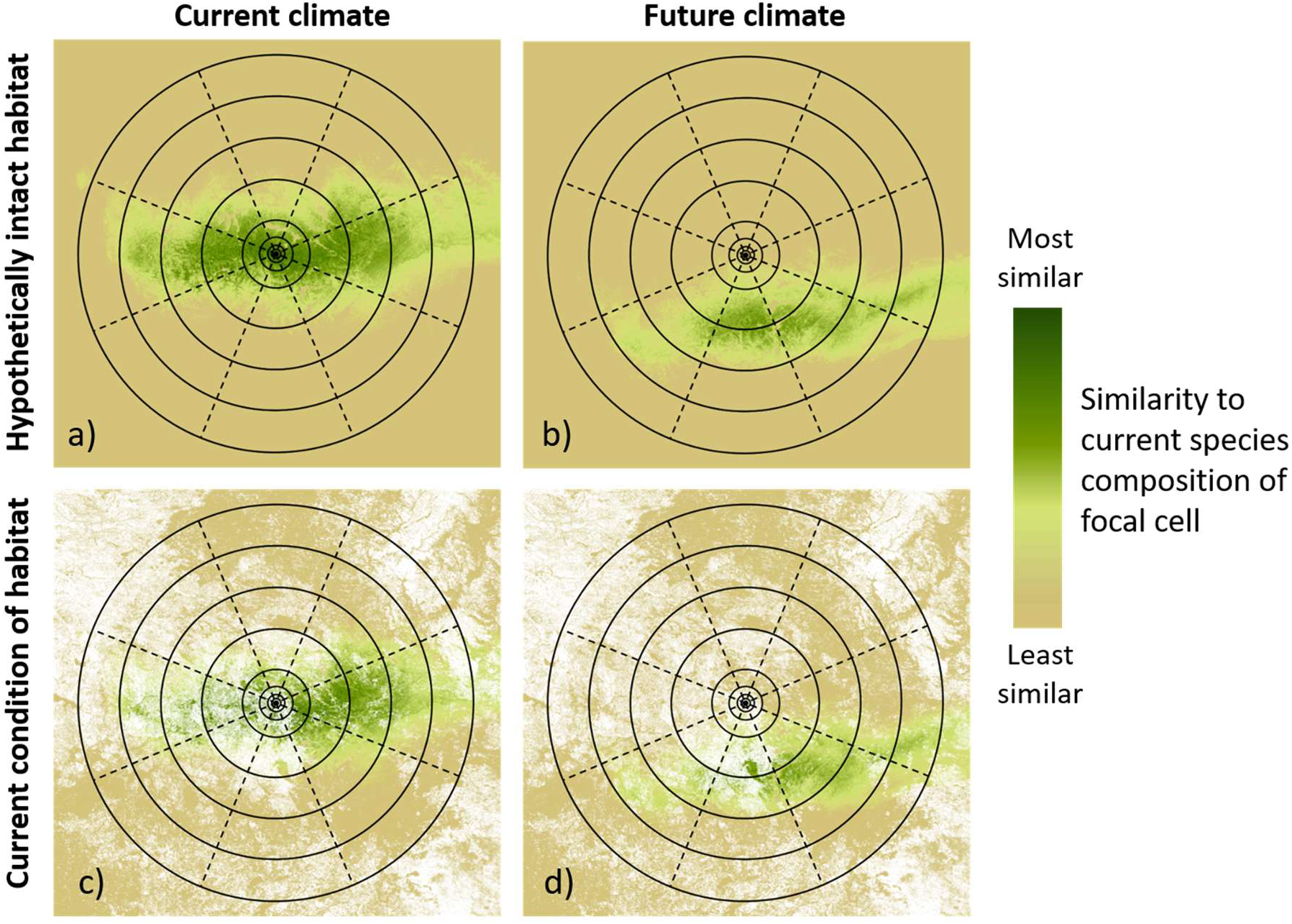
Key combinations of factors contributing to the BERI index for a given focal cell. a) Compositional similarity predicted between the focal cell and all other cells in the surrounding neighbourhood under current climate, and assuming all cells are in perfect habitat condition. b) Same as “a” but predicting similarity between the focal cell under current climate and surrounding cells under a given climate scenario. c) Same as “a” but reflecting current (observed) condition of habitat. d) Integration of “b” and “c” reflecting both current condition of habitat, and predicted shifts in compositional similarity under climate change.

For each focal cell the connectedness values calculated under the six climate scenarios described in section 2.3, along with that obtained if we assume no future change in climate (see Figure 3, panel c), are summarised into a single robust metric using the Limited Degree of Confidence (LDC) approach (Bryan et al. 2014b; McInerney et al. 2012). This is calculated as the average of: 1) the mean of connectedness values across all seven scenarios; and 2) the minimum value across the scenarios (i.e. the worst-case outcome). The BERI index is then expressed as the ratio between this summary metric of the amount of connected compositionally-similar habitat expected under climate change and *c*_*i0*_, the maximum possible value of the metric that would be obtained if the focal cell were completely surrounded by a continuous expanse of habitat in perfect condition, with no change in climate (see Figure 3, panel a):

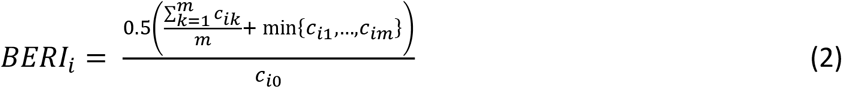

### 2.5 Sources of global data on change in habitat condition

A key input to the calculations described above is a grid of habitat condition observed at a given point in time. If, under Aichi Target 15, BERI is to be used to assess change in ecosystem resilience over time then this will further require a time-series of such grids, each reporting habitat condition for a different year of observation. Ideally these grids would satisfy three additional criteria: 1) The measure of habitat condition employed relates directly to the capacity of that habitat to support native biodiversity. 2) This measure is estimated with sufficient sensitivity, and at a sufficiently fine spatial resolution, to detect the effects both of ongoing ecosystem degradation and of ecosystem conservation or restoration actions, many of which play out over relatively small spatial extents. 3) The grids provide global coverage of all terrestrial biomes (Aichi Target 15 is not biome-specific). In reality no presently-available source of data on habitat condition adequately satisfies all of the above criteria. This situation clearly presents a challenge to operationalising the BERI indicator, but is a challenge we are currently addressing by implementing BERI in an incremental manner. This involves progressively applying the approach to ongoing improvements in spatially-explicit mapping of change in habitat condition as these products become available.

Various existing or emerging sources of data considered as candidates for initiating this process were deemed less suitable than others for this particular purpose. For example the Human Footprint dataset maps cumulative human pressures on biodiversity at relatively-fine spatial resolution across the entire terrestrial surface of the planet (Venter et al. 2016). However this product has been generated for only two time-points, 16 years apart – 1993 and 2009. While good potential now exists to derive BERI for these two points, thereby allowing an assessment of change in the index over this period, this would shed little light on shorter-term trends occurring within the 10-year period covered by the Aichi Targets (2011-2020). On a different front, the new composite indicator of land degradation being developed for Sustainable Development Goal reporting by the United Nations Convention to Combat Desertification (Sims et al. 2017), offers both high spatial and temporal resolution, but is focusing on aspects of land condition (cover, productivity, carbon stocks) not necessarily indicative of the condition of habitat for native biodiversity.

We are currently finalising development of a new global habitat-condition time series which we feel will be particularly well-suited to assessing trends in BERI. This is employing statistical downscaling of coarse-resolution land-use data using 30-arcsecond environmental and remotely-sensed land-cover covariates (Hoskins et al. 2016). We have now adapted Hoskins et al’s approach to work with Version 2, in place of Version 1, of the Land Use Harmonisation product (http://luh.umd.edu/), and with MODIS Vegetation Continuous Fields (http://glcf.umd.edu/data/vcf/) as remote-sensing covariates in place of discrete land-cover classes. Applying this downscaling approach across multiple years provides an effective means of translating observed changes in remote-sensing covariates into estimated changes in the proportions of land-use classes occurring in each 30-arcsecond cell. These proportions are then, in turn, being translated into an estimate of habitat condition, for any given cell in any given year, using coefficients derived from a global meta-analysis of land-use impacts on local biodiversity undertaken by the PREDICTS project (Newbold et al. 2016).

We further discuss this ongoing work in the final section of the paper. However, for the purposes of illustrating the derivation of BERI, and conveying the nature of results generated by this approach, we focus in this paper on an example application employing a more immediately accessible source of data on change in habitat condition – the Global Forest Change dataset http://earthenginepartners.appspot.com/science-2013-global-forest (Hansen et al. 2013). This dataset maps annual loss of forest cover from 2000 onwards, at 1-arcsecond grid resolution (approximately 30m at the equator). We use it here to generate and assess changes in BERI for a single forest biome globally – Tropical and subtropical moist broadleaf forests (here referred to simply as “Moist Tropical Forest” – as delineated by the WWF ecoregional classification (Olson et al. 2001). For any given year we estimate the condition of habitat in each 30-arcsecond cell falling within this biome (according to WWF’s mapping) as the proportion of 1-arcsecond cells it includes which are covered by forest in that year. Even though the Global Forest Change dataset also records forest gain, separately from forest loss, we focus exclusively on loss in our analysis due to challenges involved in distinguishing natural-forest gains from the establishment of exotic plantations, e.g. for palm-oil production in the tropics (Tropek et al. 2014). This means that the results presented here report only on negative changes in BERI, an issue we return to in the final section.

## 3. Results

BERI values, generated using habitat-condition data for 2014, are mapped at 30-arcsecond resolution across the Moist Tropical Forest biome in Figure 4, with a particular focus on the Indo-Malay Realm and adjacent areas. These analyses were based on the GDM models of compositional turnover for plants, and assumed a median-dispersal distance (λ in Equation 1) of 45km over the period of the scenario. This distance was selected based on broad guidance offered by Corlett and Westcott (2013) on potential rates of plant dispersal under climate change, while recognising that the time period between so-called “current” climate and 2050 scenarios in WorldClim is actually just over 60 years, given that the former represents an average of climatic observations made between 1970 and 2000.

**Figure 4:**
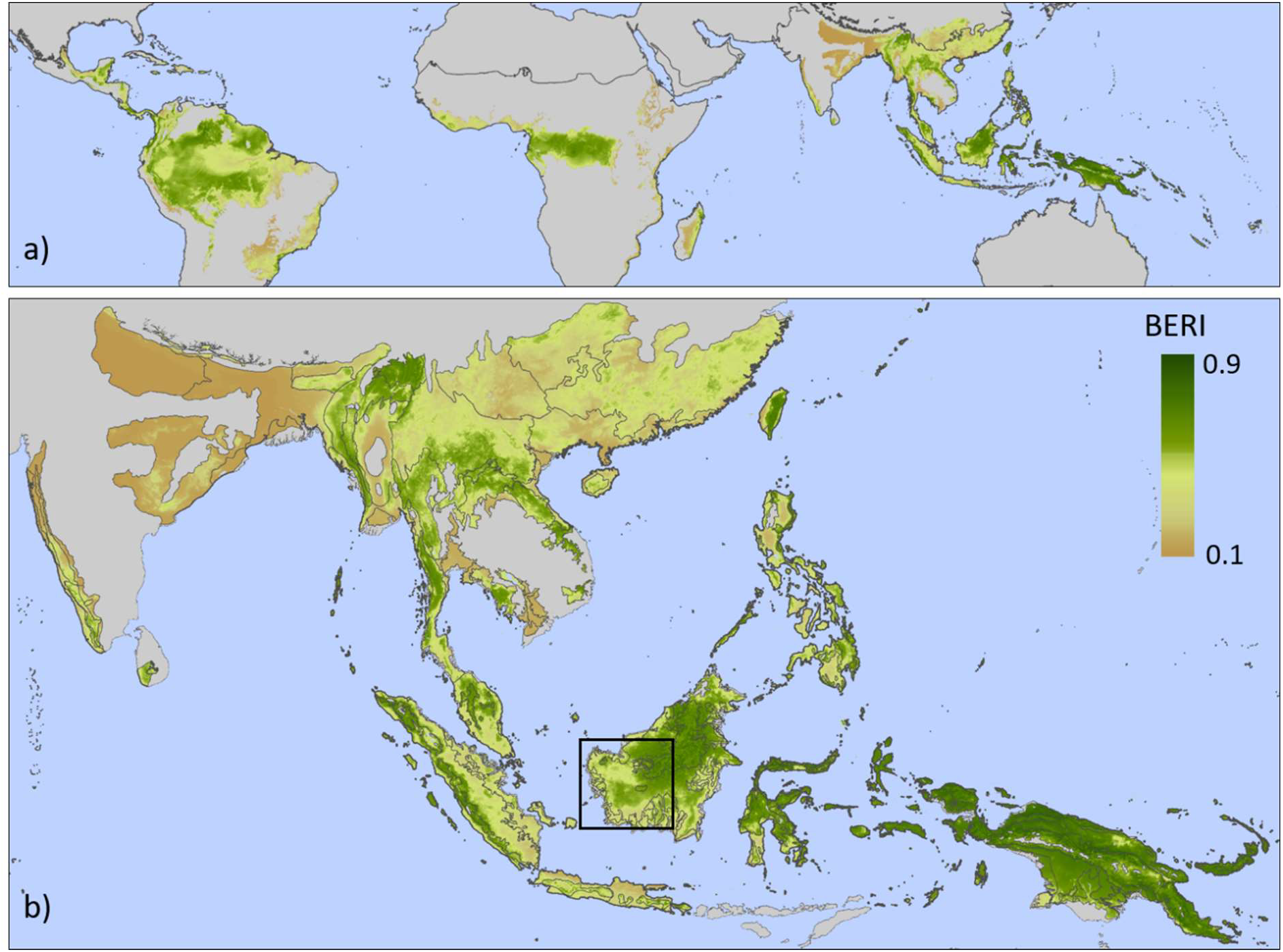
BERI mapped at 30-arcsecond grid resolution across the Moist Tropical Forest biome based on habitat condition for 2014, derived from the Global Forest Change dataset, and on compositional-turnover modelling for all vascular plants. a) Entire global distribution of the biome. b) Distribution of the biome in the Indo-Malay Realm, and adjoining portions of the Nearctic and Australasia Realms (the black rectangle delineates the area depicted in Figure 5).

Broad patterns in the spatial distribution of BERI suggest that overall levels of habitat loss and fragmentation in a landscape are likely to be key determinants of the capacity of ecosystems to retain biological diversity in the face of climate change. In general larger expanses of intact forest – e.g. in Central Africa, New Guinea, Central Borneo, and much of the Amazon – exhibit higher BERI values than less intact landscapes – e.g. in Madagascar, north-eastern India, and the Atlantic Forests of South America. However the results also suggest considerable variation in the capacity of different areas of intact forest to retain biological diversity, over and above that explained simply by the extent and condition of habitat in these areas. For example parts of the Amazon exhibit much lower values than would be expected if BERI were a function of habitat extent and condition alone. This could well be a reflection of the fact that many climate scenarios project a marked reduction in precipitation for the Amazon, as opposed to a general increase in precipitation across the tropical forests of Africa and Asia (Kooperman et al. 2018). Any such change is also likely to be further exacerbated by high levels of climate velocity associated with the flat terrain of much of the Amazon Basin – i.e. species might need to disperse quickly over long geographic distances to track changes in the distribution of their preferred climatic niche.

The complexity, and importance, of such interactions between habitat intactness and climate velocity in shaping BERI values becomes more obvious when we focus in on patterns at a regional or landscape scale. An example of this more detailed perspective is presented in Figure 5, for south-western Borneo. Comparing the 2014 habitat-condition surface (panel a) with the resulting BERI layer (panel b) for this region it is clear that the estimated capacity of forest areas to retain biological diversity under climate change is a function of more than just the extent and condition of forest habitat. To help shed more light on the interplay between relevant factors we derived two variants of BERI by experimentally assuming either: 1) that the region’s climate will undergo no change, and therefore generating the index considering purely the effects of habitat condition and configuration on retention of biological diversity (panel c); or 2) that the region is entirely covered by intact natural habitat in perfect condition, and therefore generating the index considering purely the effects of climate change on biodiversity retention (panel d).

**Figure 5:**
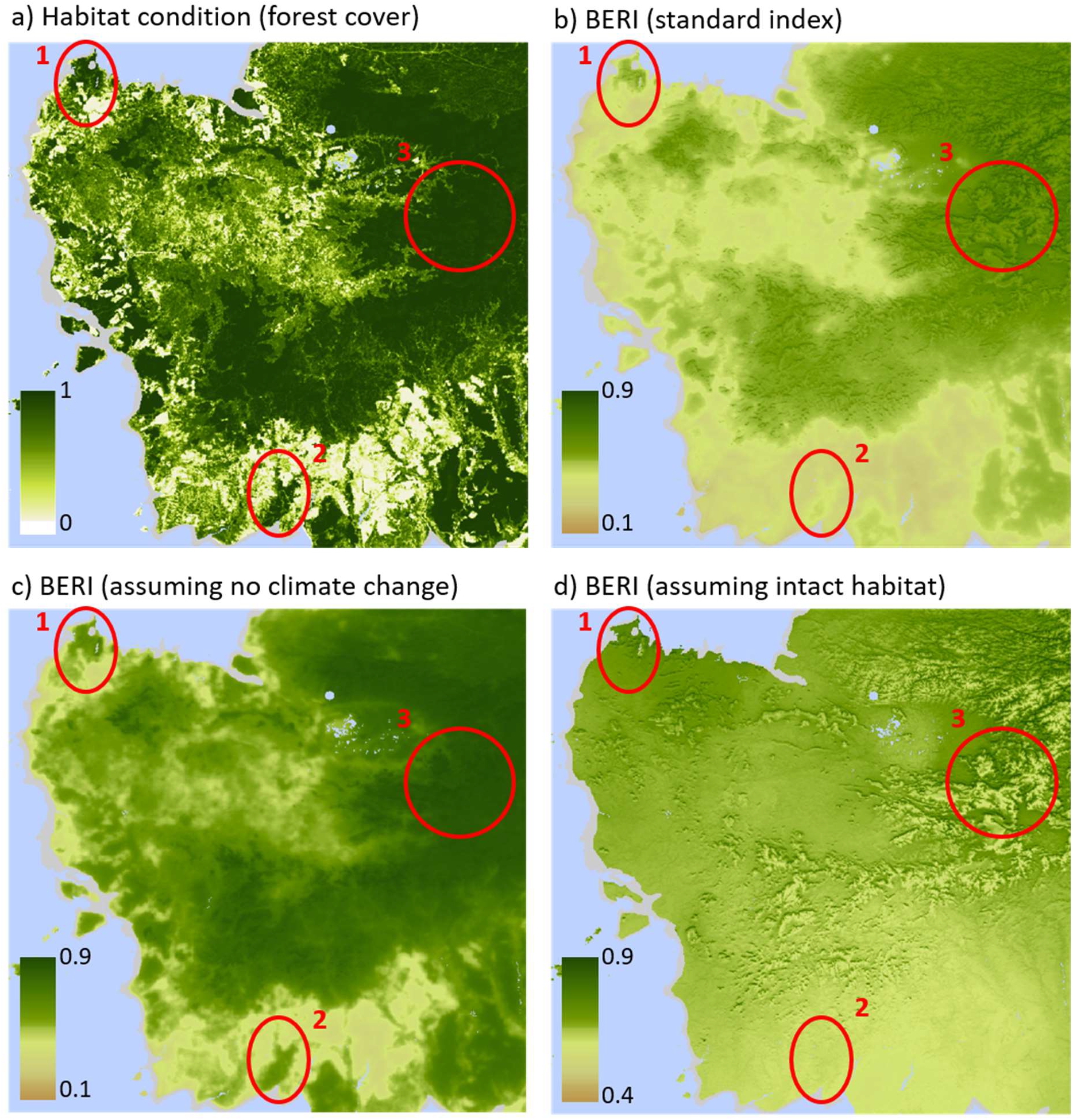
Habitat condition and BERI in 2014 for the Moist Tropical Forest biome in south-western Borneo (see Figure 4 for location map). a) Habitat condition derived from the Global Forest Change dataset, with shades of green depicting variation in the proportion of 1-arcsecond cells covered by forest within each mapped 30-arcsecond cell. b) The standard BERI index based on 2014 habitat condition, and seven climate scenarios (as also mapped in Figure 4). c) BERI based on 2014 habitat condition, but hypothetically assuming no future change in climate. d) BERI based on seven climate scenarios, but hypothetically assuming perfect habitat condition for all cells. The areas delineated with red ellipses and numbers are referred to in the text.

Highlighted areas 1 and 2 in Figure 5 contain forest exhibiting a similar level of fragmentation (panel a), yet area 1 achieves generally higher BERI scores than area 2 (panel b). The experimental variants of BERI (panels c and d) suggest that this is a function of differences in climate velocity between these two areas, most likely driven by differences in topography. Highlighted area 3 further illustrates the importance of local topography in shaping the capacity of ecosystems to retain biological diversity in the face of climate change. Within this relatively continuous expanse of natural forest, lower slopes and gullies exhibit higher BERI values than ridge and mountain tops because the present-day climatic environment of forested cells on the latter is expected to largely disappear from the landscape surrounding each of these cells.

The potential value of BERI for assessing and reporting on change in ecosystem resilience over time is illustrated in Figure 6. Here the difference in BERI resulting from observed changes in habitat condition between 2000 and 2014 is mapped for the entire Indo-Malay Realm, and trends in mean BERI values across four time points within this period are plotted for four selected ecoregions.

**Figure 6:**
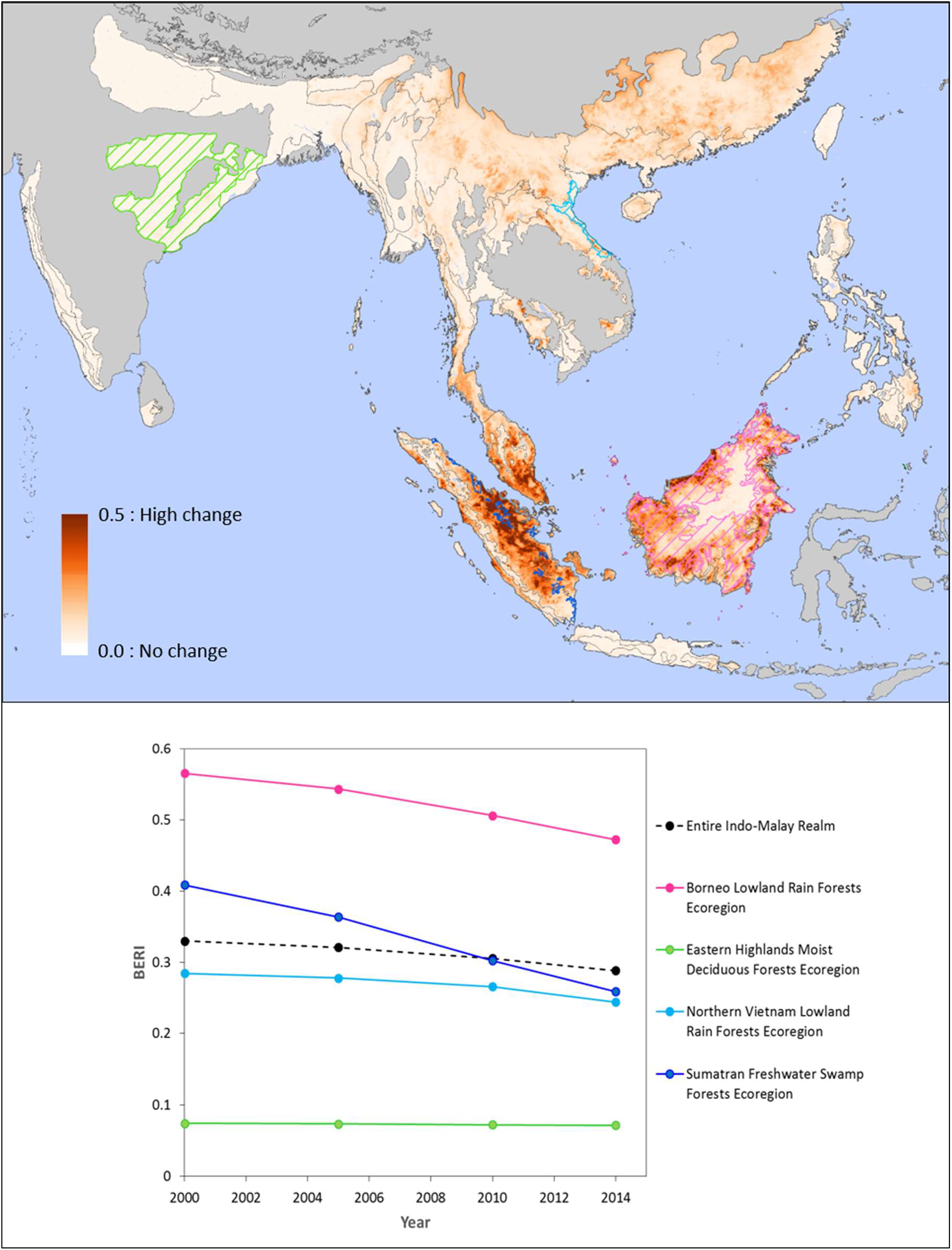
Change in BERI between 2000 and 2014 mapped for the Moist Tropical Forest biome in the Indo-Malay Realm. Estimates of change are based only on the “forest loss” component of the Global Forest Change dataset (excluding data on “forest gain”) and therefore all resulting changes in BERI are negative. The chart displays mean BERI values for the entire bio-realm, and for each of four selected ecoregions (highlighted in corresponding colours on the map).

## 4. Discussion and conclusions

The results presented here demonstrate that the BERI indicator offers considerable utility in assessing spatial variation, and temporal change, in the capacity of ecosystems to retain biological diversity in the face of climate change. The indicator effectively provides a lens through which to interpret the implications of changes in the extent and condition of natural habitat. The approach accounts not only for the gross amount of habitat remaining but also the degree to which the spatial arrangement of this habitat will allow for potential climate-induced shifts in the distribution of species. In doing so we encapsulate and integrate various existing perspectives on the roles likely to be played by climate velocity, environmental diversity (conserving nature’s stage) and habitat connectivity in determining biodiversity persistence under climate change. These concepts and principles are all involved, either explicitly or implicitly, in the derivation of BERI. Our approach complements other recent efforts to combine elements from two or more of these perspectives (e.g. Anderson et al. 2014; Belote et al. 2017; Carroll et al. 2017; Littlefield et al. 2017). However a unique strength of the method underpinning BERI is its use of compositional-turnover modelling to scale the multidimensional environmental space (including climate space) employed in our analyses.

As explained at the start of the paper, BERI has been developed primarily to enable assessment of change in a key aspect of “ecosystem resilience” under Aichi Target 15. Derivation of the index at relatively fine (30-arcsecond) spatial resolution means that results can be aggregated to report on status and trends for any desired set of reporting units – e.g. ecoregions (as in the example presented above), countries, ecosystem types, administrative or biogeographic regions, or the entire planet.

Our use of habitat-condition data derived from the Global Forest Change dataset has allowed thorough testing of the analytical techniques and software we now have in place to generate BERI from comparable input data anywhere across the world. As indicated in section 2.5 we will soon be finalising development of a habitat-condition time series covering the entire terrestrial surface of the planet at 30-arcsecond resolution. In addition to allowing BERI to be generated across all biomes (forest and non-forest) this new product will remove another significant constraint associated with the results we have presented here. Exclusion of the “forest gain” component of the Global Forest Change dataset from our analyses (see section 2.5) meant that we were limited to detecting changes in BERI resulting only from losses of natural forest, without any countering of this effect through possible improvements in habitat condition resulting from restoration efforts or natural regeneration. Because our new habitat-condition time series integrates information on both land-cover and land-use it can, at least in theory, account for both losses and gains in habitat condition, and therefore also in ecosystem resilience as assessed by BERI.

It is important to recognise, however, that any global mapping of habitat-condition change is unlikely to have sufficient sensitivity to detect restoration actions that are either highly localised (relative to the spatial resolution of this mapping), or are too recent to have yet resulted in an observable change in habitat condition. This issue deserves more attention, particularly given the strong emphasis placed on ecosystem restoration in parts of Aichi Target 15. One possible way forward might be to integrate broad-scale condition mapping with more precise delineation, at national or local scale, of areas undergoing restoration – i.e. where such detailed mapping is available use it to override that portion of the broader-scaled data when estimating BERI. While at present it appears there is no global repository of spatial data delineating areas of ecosystem restoration, it is hoped that initiatives such as the Bonn Challenge Barometer (https://infoflr.org/bonn-challenge/bonn-challenge-barometer) might help to rectify this situation in coming years.

## Acknowledgements

Development of the BERI indicator was supported by funding from the United Nations Environment Programme World Conservation Monitoring Centre (UNEP-WCMC). We sincerely thank staff at the BIP Secretariat (based at UNEP-WCMC) for their expert advice and encouragement throughout various stages of this endeavour, particularly Philip Bubb, Sarah Ivory and Anna Chenery. We also thank Rebecca Pirzl and Kristen Williams (CSIRO Land & Water) for their invaluable contribution in facilitating and supporting this project within CSIRO.

## References

Anderson, M.G., Clark, M., Sheldon, A.O., 2014. Estimating Climate Resilience for Conservation across Geophysical Settings. Conservation Biology 28, 959–970.

Beier, P., Hunter, M.L., Anderson, M., 2015. Conserving Nature’s Stage. Conservation Biology 29, 613–617.

Belote, R.T., Dietz, M.S., Jenkins, C.N., McKinley, P.S., Irwin, G.H., Fullman, T.J., Leppi, J.C., Aplet, G.H., 2017. Wild, connected, and diverse: building a more resilient system of protected areas. Ecological Applications 27, 1050–1056.

Blois, J.L., Williams, J.W., Fitzpatrick, M.C., Jackson, S.T., Ferrier, S., 2013. Space can substitute for time in predicting climate-change effects on biodiversity. Proceedings of the National Academy of Sciences of the United States of America 110, 9374–9379.

Brito-Morales, I., García Molinos, J., Schoeman, D.S., Burrows, M.T., Poloczanska, E.S., Brown, C.J., Ferrier, S., Harwood, T.D., Klein, C.J., McDonald-Madden, E., Moore, P.J., Pandolfi, J.M., Watson, J.E.M., Wenger, A.S., Richardson, A.J., in press. Climate velocity can inform conservation in a warming world. Trends in Ecology and Evolution.

Bryan, B.A., Nolan, M., Harwood, T.D., 2014a. Supply of carbon sequestration and biodiversity services from Australia’s agricultural land under global change. Global Environmental Change 28, 166–181.

Bryan, B.A., Nolan, M., Harwood, T.D., Connor, J.D., Navarro-Garcia, J., King, D., Summers, D.M., Newth, D., Cai, Y., Grigg, N., Harman, I., Crossman, N.D., Grundy, M.J., Finnigan, J.J., Ferrier, S., Williams, K.J., Wilson, K.A., Law, E.A., Hatfield-Dodds, S., 2014b. Supply of carbon sequestration and biodiversity services from Australia’s agricultural land under global change. Global Environmental Change-Human and Policy Dimensions 28, 166–181.

Buckley, L.B., Jetz, W., 2008. Linking global turnover of species and environments. Proceedings of the National Academy of Sciences of the United States of America 105, 17836–17841.

Carpenter, S., Walker, B., Anderies, J.M., Abel, N., 2001. From metaphor to measurement: Resilience of what to what? Ecosystems 4, 765–781.

Carroll, C., Roberts, D.R., Michalak, J.L., Lawler, J.J., Nielsen, S.E., Stralberg, D., Hamann, A., McRae, B.H., Wang, T.L., 2017. Scale-dependent complementarity of climatic velocity and environmental diversity for identifying priority areas for conservation under climate change. Global Change Biology 23, 4508–4520.

Corlett, R.T., Westcott, D.A., 2013. Will plant movements keep up with climate change? Trends in Ecology & Evolution 28, 482–488.

Cumming, G.S., 2011. Spatial resilience: integrating landscape ecology, resilience, and sustainability. Landscape Ecology 26, 899–909.

Drielsma, M., Ferrier, S., Manion, G., 2007. A raster-based technique for analysing habitat configuration: The cost-benefit approach. Ecological Modelling 202, 324–332.

Eigenbrod, F., Gonzalez, P., Dash, J., Steyl, I., 2015. Vulnerability of ecosystems to climate change moderated by habitat intactness. Global Change Biology 21, 275–286.

Elith, J., Leathwick, J.R., 2009. Species distribution models: ecological explanation and prediction across space and time. Annual Review of Ecology Evolution and Systematics 40, 677–697.

Ferrier, S., Harwood, T.D., Williams, K.J., 2012. Using generalised dissimilarity modelling to assess potential impacts of climate change on biodiversity composition in Australia, and on the representativeness of the National Reserve System. CSIRO, Canberra. https://research.csiro.au/climate/wp-content/uploads/sites/54/2016/03/13E_CAF-Working-Paper-13E.pdf.

Ferrier, S., Manion, G., Elith, J., Richardson, K., 2007. Using generalized dissimilarity modelling to analyse and predict patterns of beta diversity in regional biodiversity assessment. Diversity and Distributions 13, 252–264.

Fitzpatrick, M.C., Sanders, N.J., Ferrier, S., Longino, J.T., Weiser, M.D., Dunn, R., 2011. Forecasting the future of biodiversity: a test of single- and multi-species models for ants in North America. Ecography 34, 836–847.

Fitzpatrick, M.C., Sanders, N.J., Normand, S., Svenning, J.-C., Ferrier, S., Gove, A.D., Dunn, R.R., 2013. Environmental and historical imprints on beta diversity: insights from variation in rates of species turnover along gradients. Proceedings of the Royal Society B-Biological Sciences 280.

GEO BON, 2015. Global Biodiversity Change Indicators. Version 1.2. Group on Earth Observations Biodiversity Observation Network Secretariat, Leipzig. http://www.geobon.org/Downloads/brochures/2015/GBCI_Version1.2_low.pdf.

Groves, C.R., Game, E.T., Anderson, M.G., Cross, M., Enquist, C., Ferdana, Z., Girvetz, E., Gondor, A., Hall, K.R., Higgins, J., Marshall, R., Popper, K., Schill, S., Shafer, S.L., 2012. Incorporating climate change into systematic conservation planning. Biodiversity and Conservation 21, 1651–1671.

Hansen, M.C., Potapov, P.V., Moore, R., Hancher, M., Turubanova, S.A., Tyukavina, A., Thau, D., Stehman, S.V., Goetz, S.J., Loveland, T.R., Kommareddy, A., Egorov, A., Chini, L., Justice, C.O., Townshend, J.R.G., 2013. High-resolution global maps of 21st-Century forest cover change. Science 342, 850–853.

Heller, N.E., Kreitler, J., Ackerly, D.D., Weiss, S.B., Recinos, A., Branciforte, R., Flint, L.E., Flint, A.L., Micheli, E., 2015. Targeting climate diversity in conservation planning to build resilience to climate change. Ecosphere 6, 20.

Hijmans, R.J., Cameron, S.E., Parra, J.L., Jones, P.G., Jarvis, A., 2005. Very high resolution interpolated climate surfaces for global land areas. International Journal of Climatology 25, 1965–1978.

Hodgson, D., McDonald, J.L., Hosken, D.J., 2015. What do you mean, ‘resilient’? Trends in Ecology & Evolution 30, 503–506.

Hoskins, A.J., Bush, A., Gilmore, J., Harwood, T., Hudson, L.N., Ware, C., Williams, K.J., Ferrier, S., 2016. Downscaling land-use data to provide global 30’’ estimates of five land-use classes. Ecology and Evolution 6, 3040–3055.

Hoskins, A.J., Harwood, T.D., Ware, C., Williams, K.J., Perry, J.J., Ota, N., Croft, J.R., Yeates, D.K., Jetz, W., Golebiewski, M., Purvis, A., Robertson, T., Ferrier, S., 2019. Supporting global biodiversity assessment through high-resolution macroecological modelling: Methodological underpinnings of the BILBI framework. BioRxiv https://www.biorxiv.org/content/10.1101/309377v3.

Kim, H., Rosa, I.M.D., Alkemade, R., Leadley, P., Hurtt, G., Popp, A., van Vuuren, D.P., Anthoni, P., Arneth, A., Baisero, D., Caton, E., Chaplin-Kramer, R., Chini, L., De Palma, A., Di Fulvio, F., Di Marco, M., Espinoza, F., Ferrier, S., Fujimori, S., Gonzalez, R.E., Gueguen, M., Guerra, C., Harfoot, M., Harwood, T.D., Hasegawa, T., Haverd, V., Havlik, P., Hellweg, S., Hill, S.L.L., Hirata, A., Hoskins, A.J., Janse, J.H., Jetz, W., Johnson, J.A., Krause, A., Leclere, D., Martins, I.S., Matsui, T., Merow, C., Obersteiner, M., Ohashi, H., Poulter, B., Purvis, A., Quesada, B., Rondinini, C., Schipper, A.M., Sharp, R., Takahashi, K., Thuiller, W., Titeux, N., Visconti, P., Ware, C., Wolf, F., Pereira, H.M., 2018. A protocol for an intercomparison of biodiversity and ecosystem services models using harmonized land-use and climate scenarios. Geoscientific Model Development 11, 4537–4562.

Konig, C., Weigelt, P., Kreft, H., 2017. Dissecting global turnover in vascular plants. Global Ecology and Biogeography 26, 228–242.

Kooperman, G.J., Chen, Y., Hoffman, F.M., Koven, C.D., Lindsay, K., Pritchard, M.S., Swann, A.L.S., Randerson, J.T., 2018. Forest response to rising CO2 drives zonally asymmetric rainfall change over tropical land. Nature Climate Change 8, 434–440.

Littlefield, C.E., McRae, B.H., Michalak, J.L., Lawler, J.J., Carroll, C., 2017. Connecting today’s climates to future climate analogs to facilitate movement of species under climate change. Conservation Biology 31, 1397–1408.

McGuire, J.L., Lawler, J.J., McRae, B.H., Nunez, T.A., Theobald, D.M., 2016. Achieving climate connectivity in a fragmented landscape. Proceedings of the National Academy of Sciences of the United States of America 113, 7195–7200.

McInerney, D., Lempert, R., Keller, K., 2012. What are robust strategies in the face of uncertain climate threshold responses? Robust climate strategies. Climatic Change 112, 547–568.

McOwen, C.J., Ivory, S., Dixon, M.J.R., Regan, E.C., Obrecht, A., Tittensor, D.P., Teller, A., Chenery, A.M., 2016. Sufficiency and Suitability of Global Biodiversity Indicators for Monitoring Progress to 2020 Targets. Conservation Letters 9, 489–494.

Newbold, T., Hudson, L.N., Arnell, A.P., Contu, S., De Palma, A., Ferrier, S., Hill, S.L.L., Hoskins, A.J., Lysenko, I., Phillips, H.R.P., Burton, V.J., Chng, C.W.T., Emerson, S., Gao, D., Pask-Hale, G., Hutton, J., Jung, M., Sanchez-Ortiz, K., Simmons, B.I., Whitmee, S., Zhang, H., Scharlemann, J.P.W., Purvis, A., 2016. Has land use pushed terrestrial biodiversity beyond the planetary boundary? A global assessment. Science 353, 288–291.

Olson, D.M., Dinerstein, E., Wikramanayake, E.D., Burgess, N.D., Powell, G.V.N., Underwood, E.C., D’Amico, J.A., Itoua, I., Strand, H.E., Morrison, J.C., Loucks, C.J., Allnutt, T.F., Ricketts, T.H., Kura, Y., Lamoreux, J.F., Wettengel, W.W., Hedao, P., Kassem, K.R., 2001. Terrestrial ecoregions of the worlds: A new map of life on Earth. Bioscience 51, 933–938.

Pecl, G.T., Araujo, M.B., Bell, J.D., Blanchard, J., Bonebrake, T.C., Chen, I.-C., Clark, T.D., Colwell, R.K., Danielsen, F., Evengard, B., Falconi, L., Ferrier, S., Frusher, S., Garcia, R.A., Griffis, R.B., Hobday, A.J., Janion-Scheepers, C., Jarzyna, M.A., Jennings, S., Lenoir, J., Linnetved, H.I., Martin, V.Y., McCormack, P.C., McDonald, J., Mitchell, N.J., Mustonen, T., Pandolfi, J.M., Pettorelli, N., Popova, E., Robinson, S.A., Scheffers, B.R., Shaw, J.D., Sorte, C.J.B., Strugnell, J.M., Sunday, J.M., Tuanmu, M.-N., Verges, A., Villanueva, C., Wernberg, T., Wapstra, E., Williams, S.E., 2017. Biodiversity redistribution under climate change: Impacts on ecosystems and human well-being. Science 355, 1389–+.

Petrescu, A.M.R., Abad-Vinas, R., Janssens-Maenhout, G., Blujdea, V.N.B., Grassi, G., 2012. Global estimates of carbon stock changes in living forest biomass: EDGARv4.3-time series from 1990 to 2010. Biogeosciences 9, 3437–3447.

Prober, S.M., Hilbert, D.W., Ferrier, S., Dunlop, M., Gobbett, D., 2012. Combining community-level spatial modelling and expert knowledge to inform climate adaptation in temperate grassy eucalypt woodlands and related grasslands. Biodiversity and Conservation 21, 1627–1650.

Quinlan, A.E., Berbes-Blazquez, M., Haider, L.J., Peterson, G.D., 2016. Measuring and assessing resilience: broadening understanding through multiple disciplinary perspectives. Journal of Applied Ecology 53, 677–687.

SCBD, 2010. Decision adopted by the Conference of the Parties to the Convention on Biological Diversity at its tenth meeting. X/2. The Strategic Plan for Biodiversity 2011-2020 and the Aichi Biodiversity Targets. https://www.cbd.int/doc/decisions/cop-10/cop-10-dec-02-en.pdf.

Schloss, C.A., Lawler, J.J., Larson, E.R., Papendick, H.L., Case, M.J., Evans, D.M., Delap, J.H., Langdon, J.G.R., Hall, S.A., McRae, B.H., 2011. Systematic Conservation Planning in the Face of Climate Change: Bet-Hedging on the Columbia Plateau. Plos One 6, 9.

Sims, N., Green, C., Newnham, G., England, J., Held, A., Wulder, M., Herold, M., Cox, S., Huete, A., Kumar, L., Viscarra Rossel, R., Roxburgh, S., McKenzie, N., 2017. Good practice guidance. SDG Indicator 15.3.1: Proportion of land that is degraded over total land area. Version 1.0. UNCCD, Bonn, Germany.

Stein, B.A., Staudt, A., Cross, M.S., Dubois, N.S., Enquist, C., Griffis, R., Hansen, L.J., Hellmann, J.J., Lawler, J.J., Nelson, E.J., Pairis, A., 2013. Preparing for and managing change: climate adaptation for biodiversity and ecosystems. Frontiers in Ecology and the Environment 11, 502–510.

Tittensor, D.P., Walpole, M., Hill, S.L.L., Boyce, D.G., Britten, G.L., Burgess, N.D., Butchart, S.H.M., Leadley, P.W., Regan, E.C., Alkemade, R., Baumung, R., Bellard, C., Bouwman, L., Bowles-Newark, N.J., Chenery, A.M., Cheung, W.W.L., Christensen, V., Cooper, H.D., Crowther, A.R., Dixon, M.J.R., Galli, A., Gaveau, V., Gregory, R.D., Gutierrez, N.L., Hirsch, T.L., Hoft, R., Januchowski-Hartley, S.R., Karmann, M., Krug, C.B., Leverington, F.J., Loh, J., Lojenga, R.K., Malsch, K., Marques, A., Morgan, D.H.W., Mumby, P.J., Newbold, T., Noonan-Mooney, K., Pagad, S.N., Parks, B.C., Pereira, H.M., Robertson, T., Rondinini, C., Santini, L., Scharlemann, J.P.W., Schindler, S., Sumaila, U.R., Teh, L.S.L., van Kolck, J., Visconti, P., Ye, Y.M., 2014. A mid-term analysis of progress toward international biodiversity targets. Science 346, 241–244.

Tropek, R., Sedlacek, O., Beck, J., Keil, P., Musilova, Z., Simova, I., Storch, D., 2014. Comment on “High-resolution global maps of 21st-century forest cover change”. Science 344, 3.

Urban, M.C., Bocedi, G., Hendry, A.P., Mihoub, J.B., Pe’er, G., Singer, A., Bridle, J.R., Crozier, L.G., De Meester, L., Godsoe, W., Gonzalez, A., Hellmann, J.J., Holt, R.D., Huth, A., Johst, K., Krug, C.B., Leadley, P.W., Palmer, S.C.F., Pantel, J.H., Schmitz, A., Zollner, P.A., Travis, J.M.J., 2016. Improving the forecast for biodiversity under climate change. Science 353, 1113–+.

Venter, O., Sanderson, E.W., Magrach, A., Allan, J.R., Beher, J., Jones, K.R., Possingham, H.P., Laurance, W.F., Wood, P., Fekete, B.M., Levy, M.A., Watson, J.E.M., 2016. Sixteen years of change in the global terrestrial human footprint and implications for biodiversity conservation. Nature Communications 7, 11.

Walker, B.H., Salt, D., 2012. Resilience Practice: Building Capacity to Absorb Disturbance and Maintain Function. Island Press, Washington.

Williams, K.J., Prober, S.M., Harwood, T.D., Doerr, V.A.J., Jeanneret, T., Manion, G., Ferrier, S., 2015. Implications of climate change for biodiversity: a community-level modelling approach. CSIRO Land and Water Flagship, Canberra. http://adaptnrm.csiro.au/wp-content/uploads/2014/12/biodiversity-implications-tech-guide.pdf.

